# Visualization of Sympathetic Neural Innervation in Human White Adipose Tissue

**DOI:** 10.1101/2021.10.21.465246

**Authors:** Aliki Perdikari, Tessa Cacciottolo, Elana Henning, Edson Mendes de Oliveira, Julia M. Keogh, I. Sadaf Farooqi

**Author notes:** Corresponding Author: I. Sadaf Farooqi, Level 4, Wellcome-MRC Institute of Metabolic Science Box 289, Addenbrooke’s Hospital Cambridge CB2 0QQ, United Kingdom.

## Abstract

Obesity, defined as an excess of adipose tissue that adversely affects health, is a major cause of morbidity and mortality. However, understanding the structure and function of human adipose tissue has been limited by the inability to visualize cellular components due to the innate structure of adipocytes, which are characterized by large lipid droplets. Combining the iDISCO and the CUBIC protocols for whole tissue staining and optical clearing, we developed a protocol to enable immunostaining and clearing of human subcutaneous white adipose tissue (WAT) obtained from individuals with severe obesity. We were able to perform immunolabeling of sympathetic nerve terminals in whole white adipose tissue and subsequent optical clearing by eliminating lipids to render the opaque tissue completely transparent. We then used light sheet confocal microscopy to visualize innervation of human WAT from obese individuals in a 3D manner.We demonstrate the visualization of sympathetic nerve terminals in human WAT. This protocol can be modified to visualize other structures such as blood vessels involved in the development, maintenance and function of human adipose tissue in health and disease.

## 1. INTRODUCTION

Obesity-related complications such as type 2 diabetes and fatty liver disease represent a significant disease burden. As people gain weight, white adipose tissue (WAT) mass expands to store energy-rich triglycerides. Studies in rodents and in humans with inherited disorders of adipose tissue development, have shown that there is considerable variability in how much adipose tissue expansion can take place [1]. Once the adipose tissue expansion limit is reached, adipose tissue ceases to store triglycerides efficiently and lipids begin to accumulate ectopically in other tissues and organs such as the liver and skeletal muscle causing lipotoxicity, insulin resistance, cellular apoptosis and inflammation, mechanisms which underpin the development of obesity-associated metabolic complications [2].

To date, much of our understanding of adipocyte biology has emerged from studies based on histological analysis which have shown changes in adipocyte cell size, extracellular matrix (ECM) flexibility and fibrosis and macrophage infiltration in obesity [3], while the development of whole mount immunolabeling has provided insights into adipocyte precursor cell development [4]. A major function of adipose tissue in times of nutrient insufficiency is the release of stored triglycerides in WAT by lipolysis, a process which is regulated by activation of the sympathetic nervous system. However, the neural innervation of WAT has been difficult to visualize due to the presence of large cells filled with lipids [5]. Classical studies by Bartness and colleagues used retrograde tracers to perform neuronal tracing between adipose tissue and the brain in order to determine the central circuits that innervate various adipose tissue depots [6]. Recent studies in mice by Zeng and Domingos showed for the first time that white adipocytes are directly innervated by sympathetic neurons and that sympathetic innervation of WAT is necessary and sufficient to drive leptin-mediated lipolysis [7, 8]. Here we set out to optimize optical clearing techniques and whole mount immunolabeling protocols used in mice to visualize sympathetic neural circuits innervating WAT in humans.

## 2. RESULTS

### 2.1. Subcutaneous adipose tissue size from obese individuals did not alter during whole mount immunolabeling and optical clearing

We performed open biopsies of abdominal subcutaneous adipose tissue from individuals with severe obesity (Body Mass Index, BMI = 44.6 ± 5.3 kg/m^2^). Two grams of adipose tissue was immediately placed on dry ice and cut in two pieces. One piece was immersed in 4% formaldehyde (FA) and 10% sucrose in phosphate buffer saline (PBS) and kept at 4°C for 24 hours for whole mount immunolabeling, optical clearing and 3D volume imaging and the second piece was immersed in 10% neutral buffered formalin solution (NBF) and kept at room temperature for histology and haematoxylin and eosin staining (H&E) to visualize morphology in tissue sections (Figure 1A). We then optimized the iDISCO technique for whole mount immunolabeling, which is based on dehydration and rehydration of the tissue in methanol to increase adipose tissue permeability [7, 9]. After multiple rounds of organic solvent based dehydration/rehydration to remove lipids and enhance tissue permeabilization, bleaching with hydrogen peroxide to reduce tissue auto-fluorescence was performed, followed by immunolabeling with primary antibody against tyrosine hydroxylase (TH) and secondary antibody and by optical clearing with hyperhydrating solutions (Figure 1B). Although during dehydration in solutions with increased percentage of methanol the human adipose tissue biopsy shrunk, it regained its original size during the rehydration steps and the size was did not alter by the end of the dehydration/rehydration process (Supplementary Figure 1). Moreover, whilst the tissue became more rigid during the whole process it regained its original texture upon incubation for 1 hour in phosphate-buffered saline (PBS) and during the washing, permeabilization and blocking steps performed later. For immunolabeling we used an antibody against TH, which is the rate-limiting enzyme of catecholamine biosynthesis and is a marker for sympathetic nervous system. Since lipids are a major source of auto-fluorescence, which causes an increase in the background during immunolabeling for secondary antibody we used Alexa-568 fluorophore in the red spectrum, avoiding blue-green fluorophores which generate more scatter when passing through tissue.

**Figure 1.**
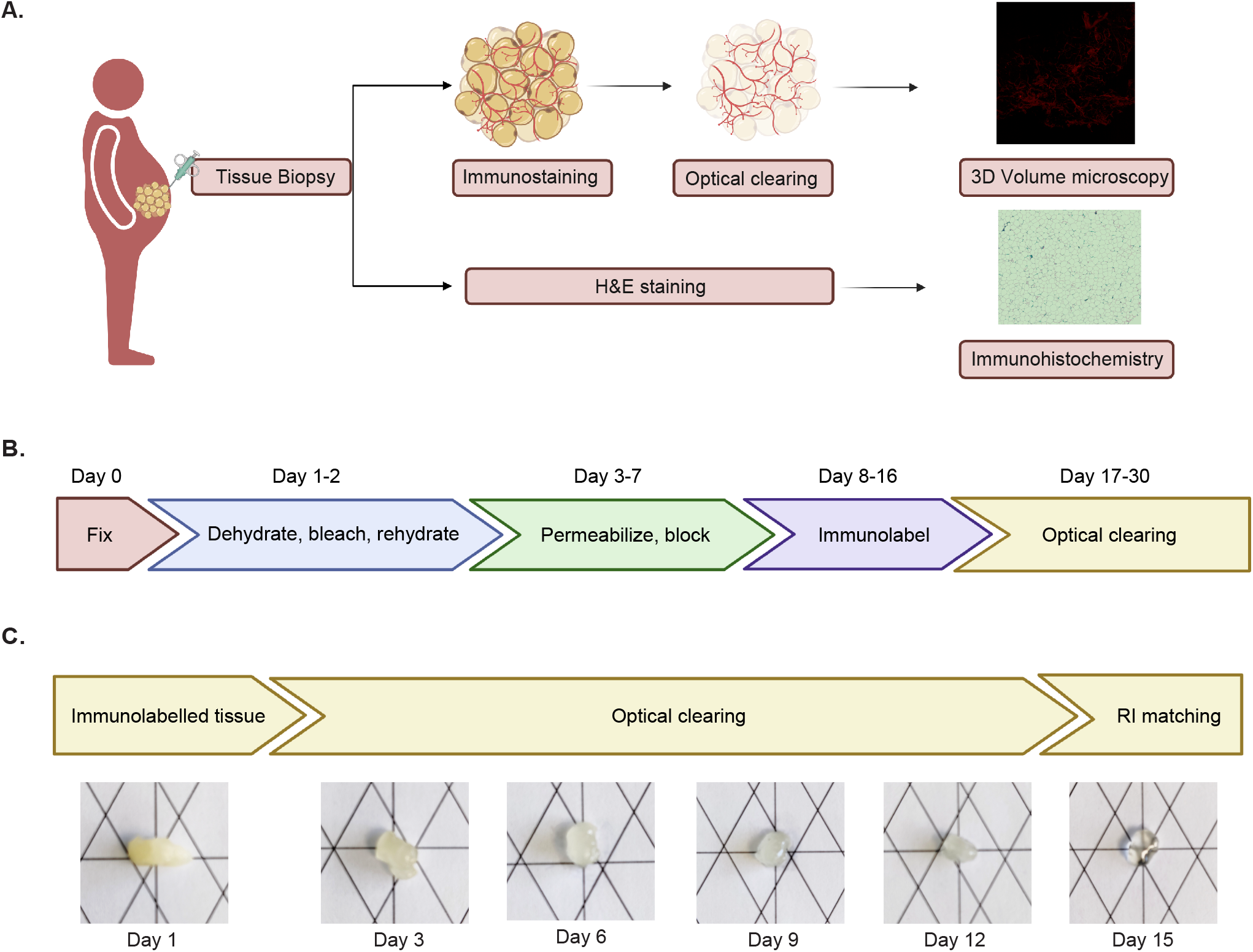
Human adipose tissue immunolabeling workflow and optical clearing. (A) Graphic representation of the use of human adipose tissue surgically removed biopsy for immunolabeling, optical clearing and 3D volume imaging and for haematoxylin and eosin (H&E) staining. (B) Graphic representation for the immunolabeling protocol for human adipose tissue. (C) Representative images for the optical clearing process of human adipose tissue after different days in ScaleCUBIC 1 and ScaleCUBIC 2 reagents.

### 2.2 Whole adipose tissue optical clearing was performed using the advanced CUBIC method

We performed adipose tissue clearing using the advanced CUBIC method, which is based on a simple immersion protocol in two different reagents to minimize light scattering [10]. Tissue was immersed in ScaleCUBIC-1 reagent, which can remove lipids, the main light scattering material inside the tissue and ScaleCUBIC-2 a reagent to further minimize light scattering and match the refractive indices (RIs) between the sample and the reagent, minimizing reflection or refraction and leading to a higher resolution image (Figure 1C). Tissue texture was altered after optical clearing, leading to a more gel like consistency due to the clearance of lipids, changing the shape of the tissue, but we did not observe any tissue expansion during or by the end of the clearing process, something documented for other tissues cleared with the CUBIC protocol [11].

### 2.3 Optical clearing with the CUBIC protocol retains adipose tissue morphology and increases whole adipose tissue light transmittance

We used H&E staining of tissue sections to confirm the presence of intact adipocytes comprised of large lipid droplets after surgically removing the human adipose tissue biopsy (Figure 2A). Moreover, we could also confirm that the adipose tissue remained intact after the biopsy by the presence of adipocytes with perilipin staining and imaging with a conventional confocal microscope (Figure 2B). After immunolabeling and optical clearing of human adipose tissue, we tested the specificity of the TH antibody to label sympathetic neural arborizations. We observed immunolabeling of neural connections in the optically cleared adipose tissue incubated with the TH antibody in comparison with tissue incubated only with the secondary antibody (Figure 2C). Furthermore, using the auto-fluorescent properties of human adipose tissue, we could show that the overall tissue architecture is remained intact even after the lengthy protocol of permeabilization, immunolabeling and clearing (Figure 2D). Lastly, we examined light transmittance in the optically cleared whole adipose tissue after using the CUBIC protocol to assess the degree of tissue clearance (Figure 2E). We could show that there is an increase of about 40% in light transmittance in the cleared sample in comparison to the light transmittance observed prior to optical clearing.

**Figure 2.**
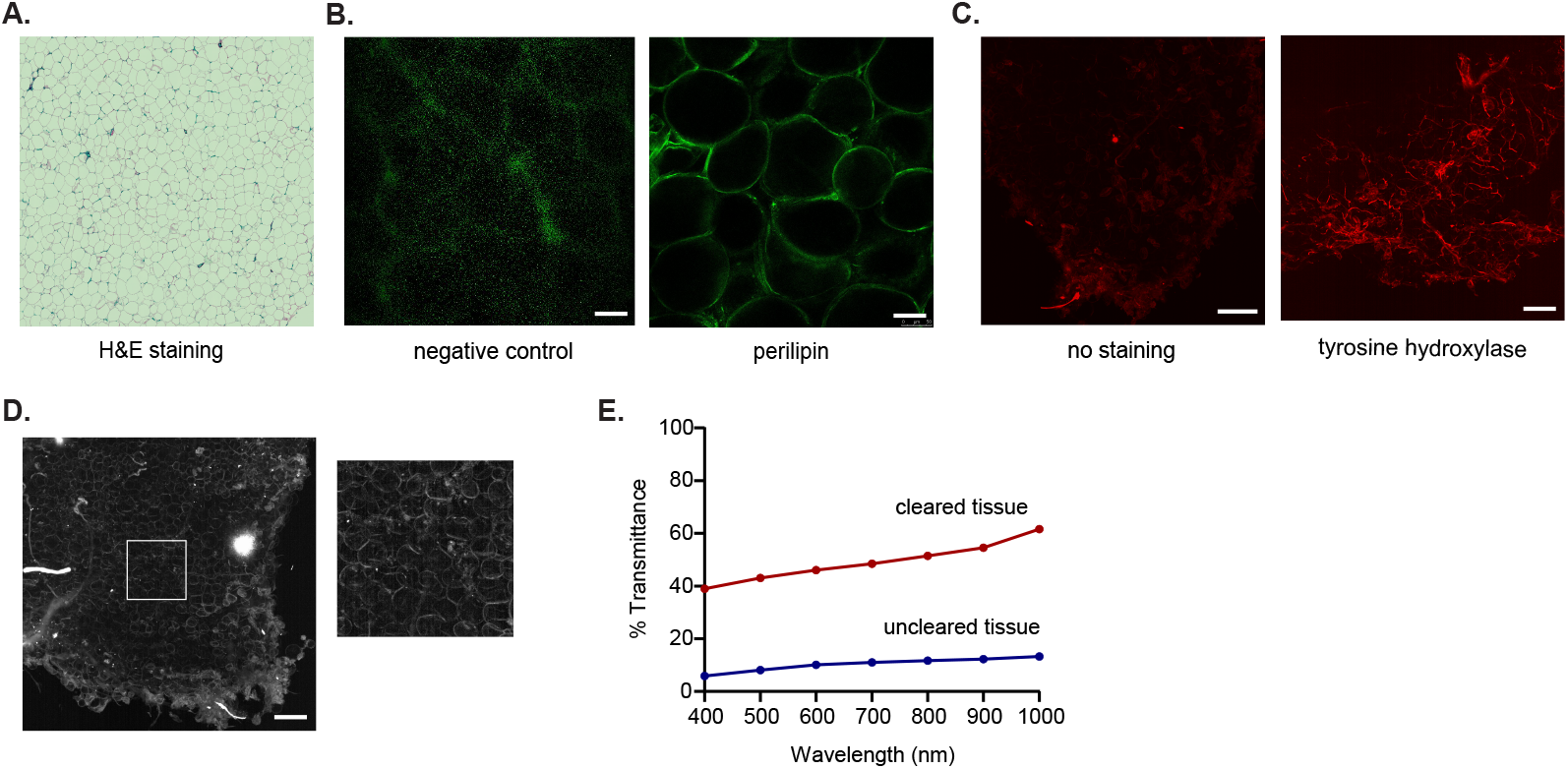
Morphology and light transmittance of human adipose tissue biopsies. (A) Representative histology image with haematoxylin and eosin (H&E) staining of human adipose tissue depicting lipid droplets of adipocytes. (B) Representative confocal image of human adipose tissue before clearing stained with perillipin or with secondary antibody only, depicting adipocytes. Green: perilipin, scale bar: 100μm (C) Maximum intensity projection images of cleared human adipose tissue without (left) or with (right) tyrosine hydroxylase staining obtained by a light sheet microscope. Red: tyrosine hydroxylase, scale bar: 200 μm. (D) Maximum intensity projection image of autofluorescence of cleared human adipose tissue, scale bar: 200 μm. (E) Quantification of light transmittance of uncleared and optically cleared adipose tissue. Data are presented as % mean of 3 individual measurements of the same tissue, normalized to blank.

### 2.4 Quantification of sympathetic innervation using whole adipose tissue 3D images

We analyzed cleared immunolabeled tissue from two severely obese people stained for TH using a light sheet confocal microscope (LSCM) that revealed in 3D the presence of sympathetic neural networks in human adipose tissue (Figure 3A-F,Movies 1 and 2). We used the 3D images obtained to calculate the density of neural arborizations (as relative number of TH+ fibers), and the mean fiber surface (Figure 2G). The fiber surface distribution revealed different size of fibers not only between the subjects analyzed but also between the different areas analyzed within the same tissue (Supplementary Figure 2).

**Figure 3.**
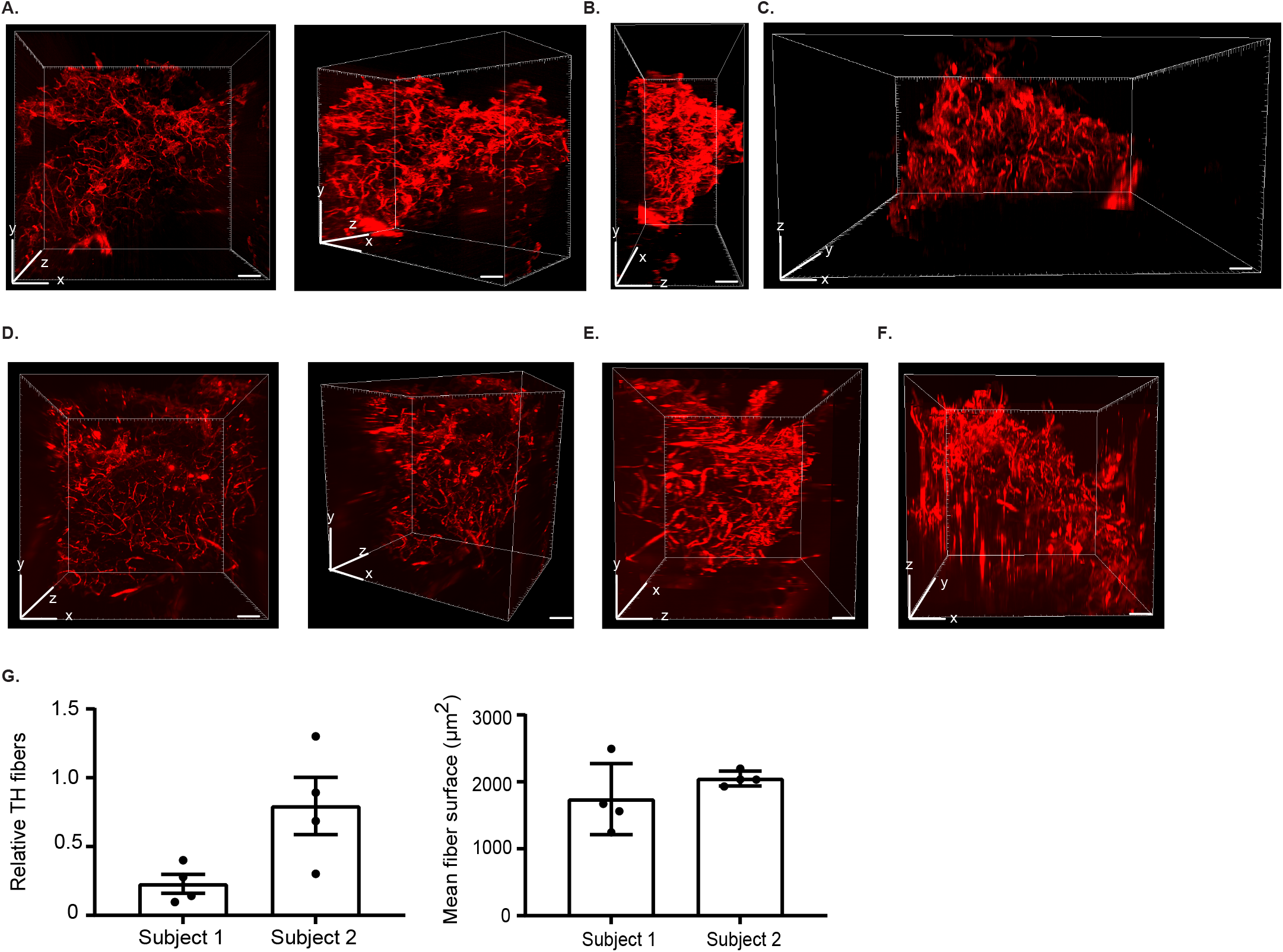
Imaging of sympathetic neural arborizations in human adipose tissue. (A-G) Representative images of the neural network in human adipose tissue obtained with light sheet microscope from two subjects [xy plane (A and D), yz plane (B and E), xz plane (C and F)], scale bar: 1mm. (G) Quantification of TH fibers presented as relative TH fibers and fiber surface, n=4 tissue areas, mean +/- standard deviation.

## 3. DISCUSSION

Until recently, studying adipose tissue innervation was limited due to conventional microscopy techniques and for several decades there has been doubt as to whether direct innervation of adipose tissue exists. The combination of tissue clearing techniques with immunolabeling and advanced 3D volume microscopy has made it possible to image whole organs and at single fiber resolution and to study sympathetic innervation in murine adipose tissue [7, 8, 12, 13]. In this study we obtained human adipose tissue biopsies from severely obese subjects and performed immunolabeling, optical clearing of the adipose tissue and 3D visualization of neural networks using a light sheet confocal microscope. Moreover, we quantified the density of tyrosine hydroxylase positive fibers in human adipose tissue samples by analyzing the 3D images obtained. Although different protocols exist for whole adipose tissue immunostaining and 3D imaging of mouse adipose tissue [7, 13], this is the first report of successful 3D imaging of sympathetic neuronal connections in human adipose tissue from severely obese subjects.

For optical clearing of human WAT, we optimized the advanced CUBIC protocol, which is based on delipidation and hyperhydration of the tissue followed by RI matching. The advantage of using the CUBIC method, despite the lengthy protocol required and a decreased clearance performance for large tissues in comparison to other clearing protocols [10], is that the solutions used are based on urea and Triton X-100 and are not hazardous, compared to the organic based solvents used in the benzyl alcohol/benzyl benzoate (BABB), AdipoClear and iDISCO protocols used for murine adipose tissue clearing, so can be easily used in any lab without special safety conditions [9, 13, 14]. Moreover, the reagents are inexpensive and technically simple to prepare and use and do not require special means for disposal. Furthermore there is no need to use any specific objective lenses for the RI match in order to image the tissue, since they are not aggressive and corrosive to the lenses of the microscope, making it possible to use different microscopes. A major disadvantage of this protocol is that clearing of the tissue takes several days that might decrease the staining intensity, while the tissue becomes highly fragile and difficult to handle in the clearing solution. To overcome these, it requires smaller sample size to minimize the time needed for optical clearing and greater attention when handling the sample to retain its structure.

There are a number of technical considerations when adapting this protocol for other uses. Prior to immunolabeling, an antibody needs to be tested for compatibility with the methanol present in the dehydration/rehydration steps of the tissue, while antibody concentration might need further adjustment. Moreover, tissue size needs to be taken into consideration when using the immunolabeling protocol and adjusted accordingly, since the size of the WAT sample will determine the time needed for effective permeabilization and blocking, as well as the length of incubation with the primary and secondary antibody. For the tissue mass used here, we required 5 days of permeabilization and blocking and 8 days of incubation in the antibody solutions.

We obtained human adipose tissue biopsies surgically, which retained the morphology of the adipose tissue but we did not try whether the same protocol could be used for adipose tissue biopsies obtained by other means, such as needle biopsies, where the needle bore and application of suction may damage tissue structure. Moreover, we obtained tissue from severely obese people, where the presence of tissue fibrosis may have impacted on immunolabeling and tissue clearing.

## CONCLUSIONS

In summary, we have developed a protocol for the optical clearing and immunolabeling of human white adipose tissue which permits the direct visualization and quantification of sympathetic innervation for the first time. Given the importance of sympathetic neural connections in mediating lipolysis and potentially other functions, the method proposed here is a powerful tool to further understand human adipose tissue structure and function and its disturbance in obesity and its metabolic complications.

## 4. MATERIAL AND METHODS

### 4.1 Study Approval

This study was approved by the Cambridge South Research Ethics Committee (18/EE/0032) and was conducted in accordance with the principles of the Declaration of Helsinki. Subjects were recruited from the Genetics of Obesity Study (GOOS) and provided written informed consent prior to inclusion to the study.

### 4.2 Subjects

Both participants had a history of severe childhood obesity, and at the time of the study had a mean age of 22.4 ± 2 years and mean BMI of 44.6 ± 5.3 kg/m^2^. They had refrained from smoking, alcohol, caffeine and strenuous exercise for 24 hours prior to the biopsy. Biopsies were carried out three hours after the consumption of a standardised lunch (35% of total energy requirements; 50% carbohydrate, 30% fat, 20% protein). Studies were performed at the Wellcome - MRC Institute of Metabolic Science Translational Research Facility

### 4.3 Surgical adipose tissue biopsy

Subcutaneous adipose tissue biopsies were obtained from the left iliac fossa, using a non-diathermy surgical biopsy method, under local anaesthesia. Participants were positioned in the supine position on a surgical couch. The skin was cleaned with 2% chlorhexidine in 70% isopropyl alcohol (Chloraprep) and covered with surgical drapes to create a sterile field. The skin was infiltrated with a maximum of 10 mls local anaesthetic (1% lidocaine with 1:200,000 adrenaline). A 2-3 cm skin incision was made, skin flaps were raised on either side of the incision and the superficial fascia incised. Adipose tissue was grasped with atraumatic forceps; around 2 grams of tissue was excised and washed in phosphate-buffered saline. Haemostasis was achieved using manual compression. A subcuticular suture (3/0 Vicryl Rapid 19 mm) was passed and reinforced with histoacryl tissue adhesive.

### 4.4 Adipose tissue fixing

For whole mount immunolabeling adipose tissue biopsy was fixed in 20ml phosphate-buffered saline (PBS) with 4% formaldehyde (Fisher Chemical) and 10% sucrose (Sigma) in a 50ml falcon tube overnight at 4°C with gentle shaking. Tissue was then washed 3 times with 5ml PBS for 1 hour each time in room temperature and then either was processed immediately for whole mount immunolabeling or kept in 70% ethanol in 4°C for maximum one month. For histology and heamatoxylin and eosin staining (H&E) adipose tissue biopsy was fixed in 10% neutral buffered formalin (NBF) in 25ml histology pots in room temperature until further processing.

### 4.5 Whole mount tissue immunolabeling

Whole mount immunolabeling protocol was adapted by Jiang et al [7]. Fixed adipose tissue was cut in 1-2cm^3^ pieces. All steps were performed in 2ml eppendorf tubes unless stated otherwise. Adipose tissue dehydration was performed at room temperature and with gentle shaking by serial incubation in 1.5ml 20% Methanol (Sigma) in double-distilled water (ddH2O) for 30minutes, 40% methanol in ddH2O for 30minutes, 60% methanol in ddH2O for 30minutes, 80% methanol in ddH2O for 30minutes and twice in 100% methanol in ddH2O for 30minutes. Dehydrated tissue was then bleached with gentle shaking at 4°C for 48 hours in 1.5 ml of 5% H2O2 (Sigma) in methanol (1 volume of 30% H2O2 in 5 volumes of 100% methanol) supplemented with 10mM EDTA and adjusted for pH 8 with Na. Tissue rehydration was performed at room temperature and with gentle shaking by serial incubation in 1.5ml 80% methanol in ddH2O for 30minutes, 60% methanol in ddH2O for 30minutes, 40% methanol in ddH2O for 30minutes, 20% methanol in ddH20 for 30minutes and twice in 1.5 ml PBS supplemented with 0.2% Triton X 100 (Fisher Chemical) for one hour. Tissue was permeabilized in 1.5ml of PBS with 0.2% Triton X 100, 20% Dimethyl sulfoxide (DMSO) (Sigma) and 0.3M Glycine (Sigma) at 37°C for 48 hours with gentle rotation and then blocked in 1.5ml of PBS with 0.2% Triton X 100, 10% DMSO and 5% donkey serum (D9663 Sigma) at 37°C for 72 hours with gentle rotation. Tissue was then immunolabeled with 1:100 anti-rabbit tyrosine hydroxylase antibody (Millipore, AB152) or 1:500 anti-goat perilipin antibody (Abcam, ab61682) in 1.5ml of PBS with 0.2% Tween20 (Sigma), 5% DMSO, 5% donkey serum and 10μg/ml heparin (Sigma) at 4°C for 96 hours with gentle rotation, washed five times in 1.5 ml of PBS with 0.2% Tween20, and 10μg/ml heparin for one hour each time with gentle rotation and labeled with the secondary antibody donkey anti rabbit Alexa 568 (Invitrogen, A-31573) or donkey anti goat Alexa 488 (Invitrogen, A-11055) in 1.5 ml of PBS with 0.2% Tween20, 5% DMSO, 5% donkey serum and 10μg/ml heparin at 4°C for 96 hours with gentle rotation. Tissue was washed 5 times in PBS with 0.2% Tween20 and 10μg/ml heparin at room temperature for 2 hours each time with gentle rotation and then kept in 1.5ml PBS at 4°C until optical clearing.

### 4.6 Tissue Optical Clearing and refractive index match

Optical clearing protocol was adapted by Susaki et al [10]. In detail, ScaleCUBIC1 (reagent 1) was prepared mixing 25 wt% final concentration Urea (Sigma), 25 wt% final concentration Quadrol 80% (Aldrich Chemical) and 15 wt% final concentration Triton X 100 in ddH20 and kept at room temperature for maximum one month. ScaleCUBIC2 (reagent 2) was prepared mixing 25 wt% final concentration Urea, 50 wt% final concentration sucrose (Sigma) and 10 wt% final concentration triethanolamine (Aldrich Chemistry) in ddH20 and kept at room temperature for maximum one month. All steps for optical clearing were performed in 2 ml eppendorf tubes. Immunolabelled adipose tissue was incubated in 1.5 ml 1:1 Reagent 1 / ddH2O solution at 37°C for 3-6hours with gentle rotation and then transferred in 1.5ml Reagent 1 solution and incubated at 37°C with gentle rotation between 8-15 days. Reagent 1 solution was refreshed every second day until tissue looked clear. After that, adipose tissue was incubated in 1.5ml 1:1 Reagent 2/PBS solution at 37°C for 3-6 hours with gentle rotation and then transferred in 1.5ml Reagent 2 solution at 37°C for 72 hours with gentle rotation until imaged in the microscope.

### 4.7 Measure of tissue light absorbance and calculation of transmittance

Uncleared and cleared adipose tissue was immersed in 100ul of CUBIC2 reagent in a well of a 96 well plate microplate with clear bottom (Greiner). Tissue absorbance measurements were taken for every 2nm between 400nm-1000nm with the Spark M10 Microplate Reader (Tecan). Blank measurements were taken from equal volume of CUBIC2 reagent in the same 96 well plates. To calculate tissue transmittance (T) from absorbance (A) we used the equation T=10^ (-A) and calculated the mean of 3 individual measurements of the same tissue. Results are presented as percentage of normalized transmittance to blank, which is set at 100%.

### 4.8 Three – dimensional volume fluorescent imaging, confocal imaging, image processing and density quantification

Optically cleared adipose tissue was imaged on a Zeiss Z1 Light sheet microscope (Zeiss) in clearing configuration using a single or a dual-side illumination, with the EC Plan-Neofluar 5x/0.16 Air immersion objective, a 561nm laser illumination and the BP 575-615nm emission filter. Tissue was attached with glue in the edge of a syringe in order to be immobilized and immersed in the imaging chamber filled with ScaleCUBIC 2 (Reagent 2) solution carefully to avoid bubble formation. Acquisition and initial image processing was performed using the ZEN black software (ZEISS,2014 service pack 1 version 9.2.6.54), stitching was performed in Arivis software (Arivis) and further image processing in Fiji and ZEN lite (ZEISS). Imaging of perilipin in uncleared adipose tissue was performed using a Leica SP8 confocal microscope (Leica Microsystems).

Quantification of tyrosine hydroxylase fiber density and fiber surface (μm^2^) was performed using the Arivis Vision4D software (Arivis). Fiber volume was calculated (μm^3^) in 4 Z-stacks areas per subject using the fluorescent signal and the blob function. Fibers with volume < 600 μm^3^ were not included in the analysis. Results were calculated by fiber volume (μm^3^) divided to total area volume (μm^3^) and presented as mean relative TH fibers. Tissue surface area (μm^2^) was calculated with Imagej (Fiji). Graphs were prepared with Prism 7 (Graph Pad Software).

## AUTHOR CONTRIBUTIONS

Aliki Perdikari: conceptualization, formal analysis, investigation, writing-original draft preparation, visualization; Tessa Cacciottolo: resources, writing-original draft preparation; Elana Henning: resources, writing-original draft preparation; Edson Mendes de Oliveira: investigation, writing-original draft preparation; Julia M. Keogh: resources, writing-original draft prepapartion, project administration; I Sadaf Farooqi: conceptualization, writing-original draft preparation, resources, project administration.

## ACKNOWLEDGEMENTS

We thank the patients and families for their involvement. AP was supported by the Swiss National Science Foundation (P400PB_186783). I.S.F. was supported by the Wellcome Trust (207462/Z/17/Z), the NIHR (National Institute for Health Research) Cambridge Biomedical Research Centre, Botnar Fondation, the Bernard Wolfe Health Neuroscience Endowment and a NIHR Senior Investigator Award. Clinical studies were performed in the Wellcome-MRC IMS Translational Research Facility (TRF) supported by a Wellcome Trust Major Award (208363/Z/17/Z). Light sheet confocal microscopy was performed in the Gurdon Institute Imaging Facility. Histology was performed in the Wellcome-MRC IMS Histopathology Core Facility and confocal and histology imaging was performed in the Wellcome-MRC IMS Image Core Facility. We would like to thank Nicola Lawrence and Alex Sossick at the Gurdon Institute Imaging Facility for microscopy and image analysis support. We would also like to thank Sarah Kohnke from Clemence Blouet lab in Wellcome-MRC IMS for support with the clearing protocol.

The funding bodies had no role in the design or conduct of the study; collection, management, analysis or interpretation of the data; preparation, review or approval of the manuscript or the decision to submit the manuscript for publication.

## FIGURE LEGENDS

**Supplementary Figure 1. Human adipose tissue optical clearing prior to whole mount immunolabeling**. Representative images of human adipose tissue after the dehydration and rehydration steps of immunolabeling.

**Supplementary Figure 2. Distribution of fiber surface per area analyzed**. Fiber size is depicted as fiber surface (μm^2^) calculated automatically by the Arivis Vision4D software. Each graph depicts the distribution of fiber surface in the area analyzed.

**Movie 1 – Supplementary to Figure 3**. 3D visualization of human adipose tissue neural innervation in one region of interest from the sample analyzed.

**Movie 2 –Supplementary to Figure 3**. 3D visualization of human adipose tissue neural innervation in the whole sample analyzed, performed by stitching different regions of interest.

