## Supplementary figures and images for "Visualization of Sympathetic Neural Innervation in Human White Adipose Tissue"

### Supplementary Figure 1

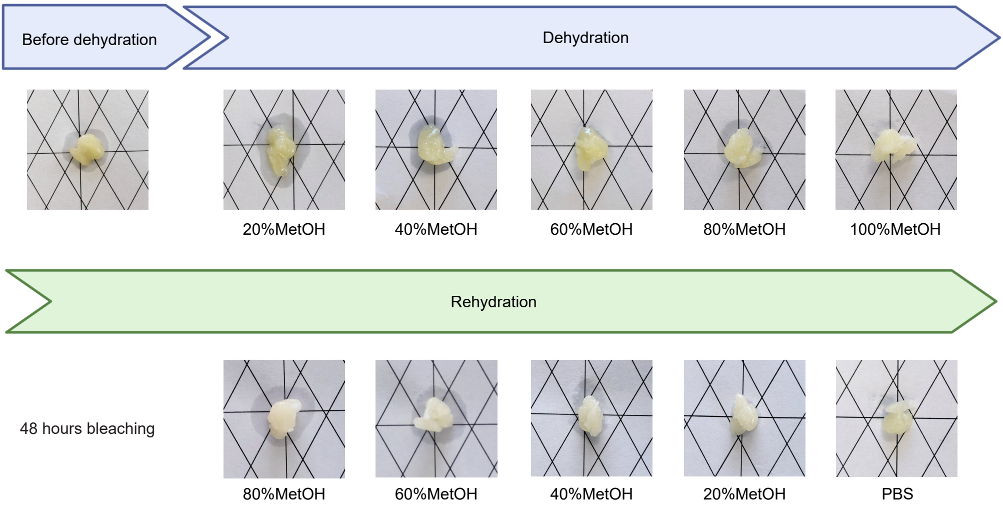

### Supplementary Figure 2

**Subject 1**

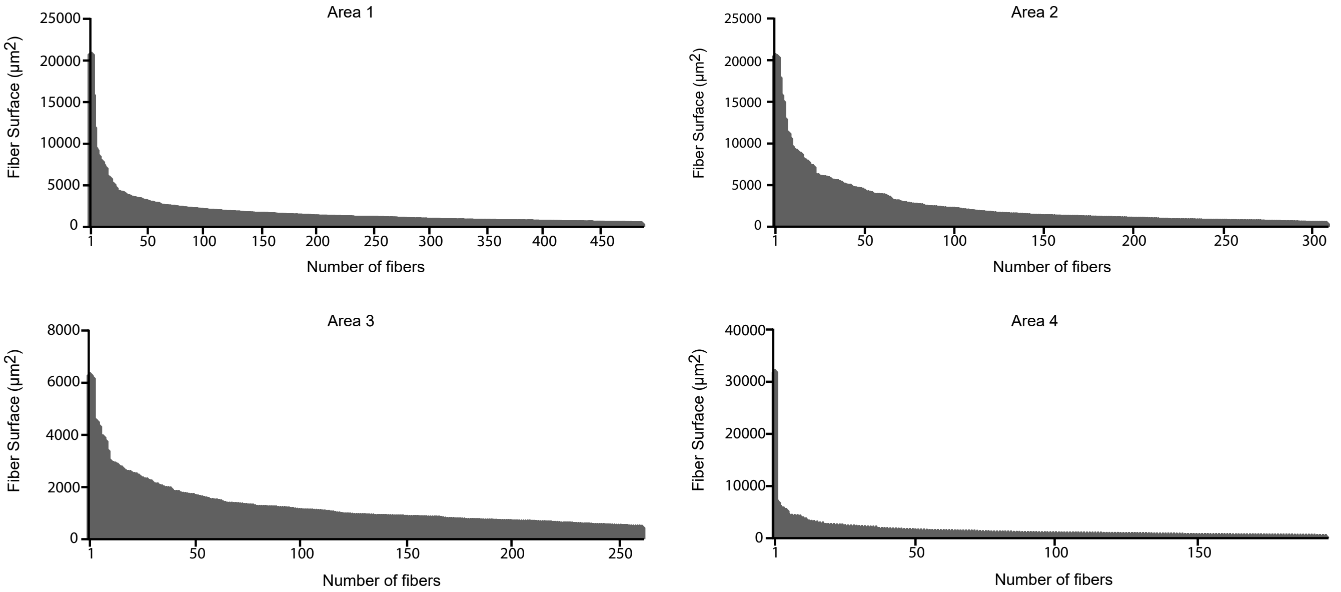


**Subject 2**

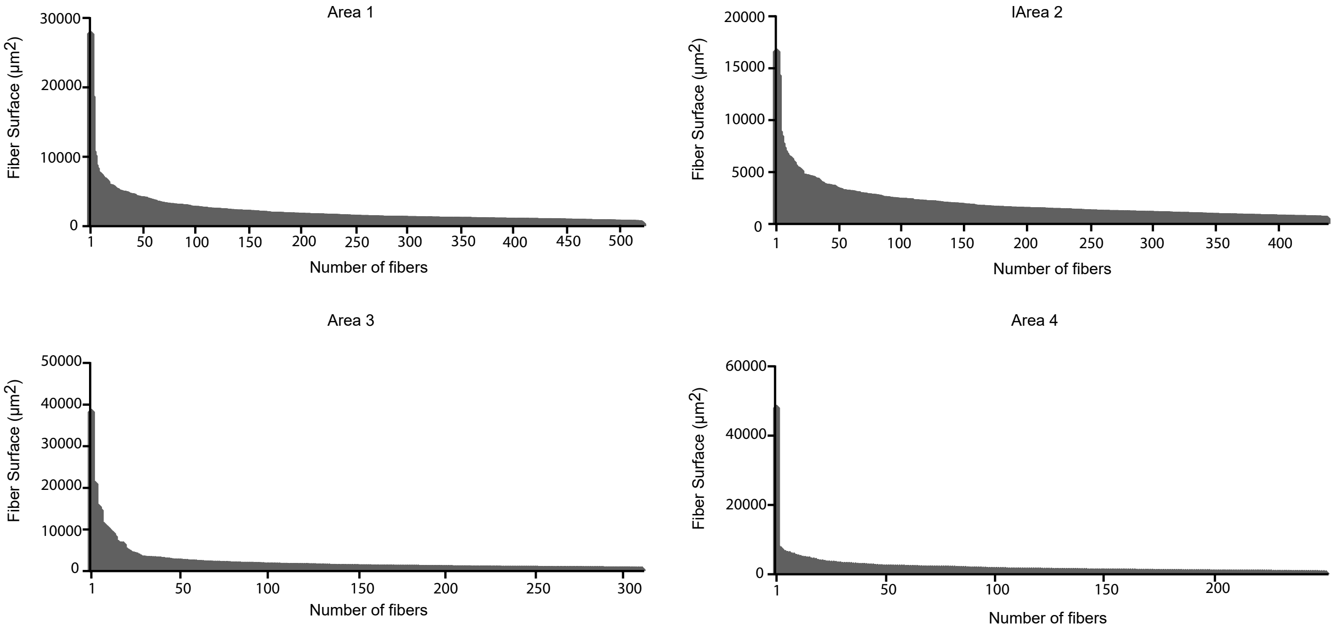
